# A new paradigm for personalized cancer screening

**DOI:** 10.1101/265959

**Authors:** Rachel Walker, Heiko Enderling

## Abstract

A series of distinct histologic lesions precedes the onset of malignancy in many common cancers, yet early detection remains a major challenge. Many patients still experience late (stage IV) diagnoses and thus poor prognosis and limited options for therapeutic intervention. For cancers with known biomarkers of premalignant progression, optimized patient-specific screening protocols would minimize the risk of undetected progression to advanced stage disease. Here, we propose simple, cost-effective mathematical and statistical approaches to forecasting disease progression that could guide the personalization of optimal screening times for high-risk patients.

## I. INTRODUCTION

In many common cancers the onset of malignancy is not an instantaneous event but rather the gradual, progressive development of advancing precursor lesions. Such pre-cancerous conditions are often used to identify patients at high risk of eventual malignancy. In the stomach and esophagus, prolonged tissue damage induced by *Helicobacter pylori* infection or gastroesophageal reflux disease, respectively, can initiate a pathway from metaplasia to dysplasia and ultimately carcinoma^1,2^. Ulcerative colitis and crohn disease can develop into colon cancer^3^, and chronic hepatitis B virus infection is the most common cause of liver disease, causing fibrosis, cirrhosis and eventually hepatocellular carcinoma^4^. Despite this, it is not always necessary, possible or cost-effective to intervene clinically at these early stages as a preventative measure.

Only a small fraction of patients ultimately progress to advanced stage disease, and progression times through carcinogenesis can be highly patient-specific. After establishing which patients are at high risk, ideally we would efficiently monitor this population to minimize the chance of undetected malignant transformation. There is unutilized potential for established biomarkers and simple mathematical and statistical tools to guide the efficient monitoring of disease development for early detection on a patient-by-patient basis.

Many upregulated proteins and differentially expressed genes have been identified in many cancer types^5^. These biomarkers can typically be quantified both in healthy patients and in the presence of malignancy through either tissue biopsy or blood sample, allowing the identification of expression ranges most frequently associated with disease and the subsequent prospective determination of if a patient is at “high risk”. From an early detection perspective, a clinically useful output may be the patient-specific rate of change of these biomarkers, providing insights into disease progression rates and thus facilitating recommendations for optimal, patient-specific screening scheduling.

Simple and cost-effective techniques are readily available to quantify the dynamic expression behavior of these biomarkers using as few as two clinical measurements. With particular emphasis on cancers in which patients are frequently diagnosed with an early, precancerous condition, evaluation of biomarker expression at initial presentation and in subsequent follow-up may be the only information required to predict the future trajectory of biomarker expression and corresponding disease progression in an individual patient.

## II. H.PYLORI(+) GASTRIC CANCER AS AN ILLUSTRATIVE EXAMPLE

The primary cause of gastric cancer worldwide is infection with *Helicobacter pylori*, which initiates a pathway of advancing histologic lesions from gastritis to intestinal metaplasia, dysplasia and cancer^1^. At present, many patients initially present with *H.pylori*-associated gastritis and receive antibacterial treatment. It is now understood that bacterial eradication does not eliminate the risk of disease progression^6-8^, yet no follow-up screening protocol exists for high-risk patients with a history of the bacterial infection. Many of these patients ultimately return to the clinic with untreatable, stage IV disease. Progression time varies dramatically between individuals. This necessitates patient-specific, personalized follow-up screening protocols to reduce the high mortality rate of this disease.

Several biomarkers including CD44, CD133 and Lgr5 have been found to be upregulated in gastric cancer tissue^9,10^. Immunohistochemical staining of tissue samples from each respective stage of this pathway demonstrated that expression of these biomarkers increased in a step-wise fashion throughout carcinogenesis^11^. This presents an opportunity to utilize these known biomarkers to predict disease progression for future patients presenting at early stages of the carcinogenic pathway.

### A. Mathematical Approach

There are several available approaches to quantifying the patient-specific dynamics of biomarker positivity. The first requires only a simple ordinary differential equation (ODE) describing the rate of change of marker expression (M) in time (dM/dt). Two such examples (linear growth model, exponential growth model) are demonstrated in Figure 1. Each equation features one parameter, the rate of change of marker expression, α, that is highly patient-specific. Simple data fitting techniques^12^ can provide this value for any patient for whom two or more clinical values are available. Figure 2 demonstrates the significant variation in this parameter between individual gastric cancer patients who all progressed to adenocarcinoma at different times, and thus experienced varying growth rates α of the marker-positive population.

**Figure 1.**
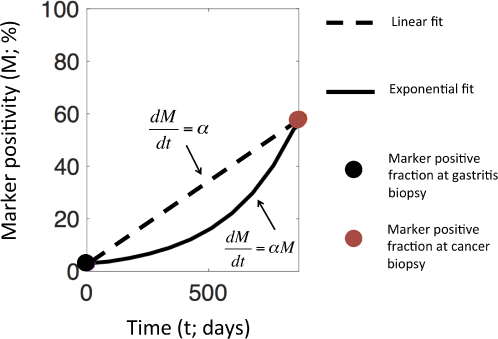
CD44 marker-positivity of an individual patient from immunohistochemical staining of routine tissue samples at date of gastritis diagnosis (black circle) and date of cancer diagnosis (red circle). Fitting linear (dashed line) and exponential (solid line) growth models to these points allows growth rate quantification.

**Figure 2.**
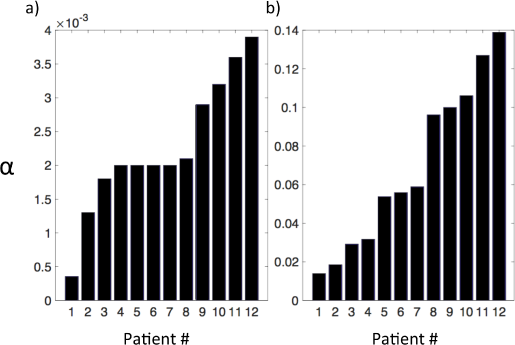
Values of patient-specific growth rate of marker positivity (α) for 12 *H.pylori* (+) gastric cancer patients from a cohort in Cali, Colombia^11^. Values obtained from fitting linear model (left panel) and exponential model (right panel) to sequential biopsies of each patient.

Depending on the specific biological problem, such simple one-term growth models can be expanded to more complex systems of ordinary differential equations based on our understanding of the underlying biology, with additional terms added to represent specific mechanisms of increase. For example, CD44, CD133 and Lgr5 are known biomarkers of stem cells; the increased proliferation of stem cells at the site of inflammation or recruitment of stem cells from the bone marrow to contribute to tissue repair have been incorporated in models of patient-specific increase in stem cell fraction during gastric carcinogenesis^13^.

If patient-specific biomarker increase rates are routinely quantified in this manner, they can be utilized for disease progression predictions in future patients. Potential trajectories of the marker-positive population can be simulated based on the clinically observed behavior of past patients coupled with the initial condition of the new patient at presentation^14^. Based on the historical data, probability distributions can be mapped onto all potential outcomes to identify the most common clinically observed progression pathways (Figure 3a). This can guide the recommendation of an appropriate follow-up screening time on a patient-by-patient basis. After only one follow-up, the actual progression rate of a patient can be calculated and used to predict the future trajectory of biomarker increase and corresponding disease progression using a Bayesian adaptive prediction approach^15^ (Figure 3b). As the number of measurements increases, the accuracy of this estimation will simultaneously increase. The overall goal of such approach would be to detect transition to malignancy at the earliest possible opportunity, allowing therapeutic intervention at a time that would maximize the likelihood of success.

**Figure 3.**
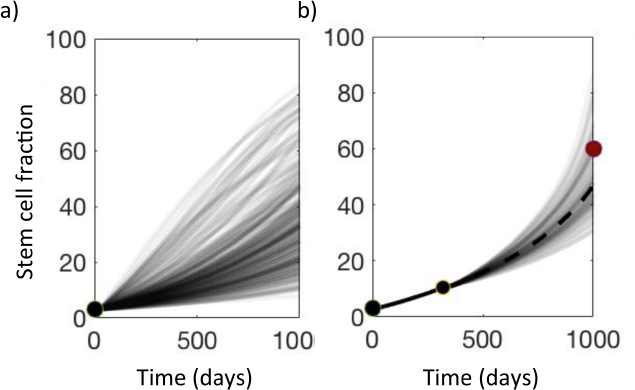
**a)** Potential growth trajectories of the marker-positive population from a known initial condition in gastritis (black circle). Darker lines represent higher likelihood of outcome. **b)** With only two clinically obtained data points (black circles), patient-specific growth rate α can be calculated, and future growth trajectory simulated. Where marker expression is known to correlate with disease progression, this can allow the recommendation of follow-up screening times to minimize the risk of undetected malignant transformation.

### B. Statistical Approach

To maintain mechanism-independence, or if mechanisms are unknown, basic statistical tools could be utilized to recommend optimal screening times on a patient-by-patient basis. Using simple logistic regression approaches^16^, the likelihood of a patient with certain characteristics (e.g age, gender, current biomarker-positivity) being at an advanced stage of disease can be calculated based on historical data from clinical patients. Coupling these likelihoods with the average growth rate of marker-positivity facilitates the calculation of an expected time until this advanced stage is reached for prospective clinical patients, based on their current relevant characteristics. We can then not only stratify patients according to their current risk status, but also suggest follow-up screening times prior to the predicted onset of advanced stage disease. Application of this approach in the gastric cancer setting is described in [11].

## III. DISCUSSION

In many common cancers, biomarkers of carcinogenesis are well established. Such markers are commonly used to identify patients at high risk, but are rarely utilized for improving screening. With only two sequential clinical observations it is possible to quantify the growth rate of an individual patients’ biomarker expression and predict how this expression will develop in the future. The blood, serum or biopsy data necessary to conduct this simple analysis can be obtained during early routine clinical visits. Computationally, the identification of patient-specific parameters and simulation of future progression can be completed almost instantaneously on one of many readily available software packages such as MATLAB.

At present, methods for screening scheduling are generic and often outdated. In several cancers including that of the stomach, follow-up screening after diagnosis with precursor lesions such as gastritis is not even required clinically despite many patients continuing to advanced stage disease. In this era of precision medicine we need to extend our efforts to also personalize prevention strategies. A discussion needs to be initiated on how known biomarkers of carcinogenesis can be coupled with simple and readily available quantitative tools to improve screening protocols and reduce the currently vast number of cases of late-diagnosis induced mortality.

